# Altered resting state functional connectivity in a thalamo-cortico-cerebellar network in patients with schizophrenia

**DOI:** 10.1101/506576

**Authors:** Caroline Garcia Forlim, Leonie Klock, Johanna Bächle, Laura Stoll, Patrick Giemsa, Marie Fuchs, Nikola Schoofs, Christiane Montag, Jürgen Gallinat, Simone Kühn

## Abstract

Schizophrenia is described as a disease in which complex psychopathology together with cognitive and behavioral impairments are related to widely disrupted brain circuitry causing a failure in coordinating information across multiple brain sites. This led to the hypothesis of schizophrenia as a network disease e.g. in the cognitive dysmetria model and the dysconnectivity theory. Nevertheless, there is no consensus regarding localized mechanisms, namely dysfunction of certain networks underlying the multifaceted symptomatology.

In this study, we investigated potential functional disruptions in 35 schizophrenic patients and 41controls using complex cerebral network analysis, namely network-based statistic (NBS) and graph theory in resting state fMRI. NBS can reveal locally impaired subnetworks whereas graph analysis characterizes whole brain network topology.

Using NBS we observed a local hyperconnected thalamo-cortico-cerebellar subnetwork in the schizophrenia group. Furthermore, nodal graph measures retrieved from the thalamo-cortico-cerebellar subnetwork revealed that the total number of connections from/to (degree) of the thalamus is higher in patients with schizophrenia. Interestingly, graph analysis on the whole brain functional networks did not reveal group differences. Together, our results suggest that disruptions in the brain networks of schizophrenia patients are situated at the local level of the hyperconnected thalamo-cortico-cerebellar rather than globally spread in brain.

Our results provide further evidence for the importance of the thalamus and cerebellum in schizophrenia and to the notion that schizophrenia is a network disease in line with the dysconnectivity theory and cognitive dysmetria model.

## 1. Introduction

Schizophrenia is a mental disorder that includes alterations in cognitive processes, perception, affect, and the sense of self, with symptoms spanning from so-called positive symptoms (hallucinations and delusions) to negative symptoms (alogia, anhedonia, apathy, and attentional impairments)^1^. Within a lifetime, around 7 out of 1,000 individuals develop schizophrenia and in those the risk of dying is two to three times increased compared to the general population emphasizing the importance of this disorder to the public health^2,3^. As a comprehensive theory of schizophrenia should be capable of explaining the range of heterogeneous symptoms associated with the illness^4^, the variety of symptoms constitute a central challenge to schizophrenia research^5^. From a neurophysiological perspective, the research focus of schizophrenia has moved from investigating impaired localized brain regions and their relationship with psychopathology to studying abnormal interactions between brain regions^5^ giving rise to models of altered neural connectivity that claim to explain the heterogeneity of symptoms distinctive for schizophrenia. Prominent examples are the cognitive dysmetria and disconnection hypothesis^4–6^). Indeed, within the last years a growing body of neuroimaging studies report aberrant brain circuitry causing a failure in coordinating information across multiple brain sites^7–11^.

Employing magnetic resonance imaging (MRI) allows an exploration of the connectivity across the entire brain. The organization of the brain can be represented by a network that is comprised of nodes (close or distant cortical brain areas) that are connected via links (functional or structural correlations) with each other. From this perspective, functional and structural organization of the human brain can be investigated and by that information processing among disparate brain regions^12^. The connectivity of the brain can be studied when individuals engage in specific cognitive processes elicited by a certain task or when they are lying awake without such an external attention focus in the scanner, the latter is called resting state. The aim of resting-state functional MRI (rsfMRI) is to measure the intrinsic or “default” brain activity, the activity that is associated with almost the totality of the brain energy consumption, nearly 95%^13^. In rsfMRI, functional brain networks are often constructed based on brain areas that were obtained from anatomical templates for nodes and temporal correlations given by Pearson’s correlation coefficient for links.

Once the functional networks are built, the next step is to compare the networks. In order to find potential differences between groups of individuals, the network comparison is usually conducted pairwise between all links. This pairwise comparison leads to a large number of group comparisons and therefore, multiple comparison problems. To overcome this issue and yet controlling for family-wise error, an approach called network-based statistic (NBS) has been developed^14^ NBS is a nonparametric analysis rooted on cluster-based thresholding of statistical parametric maps and thus is able to identify local subnetwork differences between groups.

Another widely used method to compare functional brain networks is graph analysis where information processing can be characterized by means of network topology. The network topology is obtained by employing graph measures that fall into groups that are associated with different aspects of information processing: centrality, functional integration, and functional segregation. Centrality measures the importance of a node in the network and is related to the concept of a hub. Functional segregation is associated with clustering and therefore with specialized local information processing. Functional integration relies on the concept of paths/distances and is associated to the global exchange of information in the brain. In contrast to NBS, the networks are built first, then topological properties are calculated and as a last step, group comparisons are performed.

In the present study, we aimed to investigate resting-state functional connectivity in a group of patients diagnosed with schizophrenia and a group of matched healthy individuals. Therefore, we employed 2 complimentary methods, NBS to identify potentially impaired subnetworks among groups and graph analysis, to characterize the topology of the functional networks and examine potential topological differences.

## 2. Methods

### Participants

Forty-one healthy individuals and thirty-five individuals who met the criteria for a diagnosis of schizophrenia following the International Classification for Diseases and Related Health Problems (ICD-10) were included in the reported analysis. Patients were recruited at St. Hedwig Hospital of the Charité-Universitätsmedizin Berlin (Germany). The severity of symptoms was rated by a trained clinician with the Positive and Negative Syndrom Scale (PANSS)^15^ of which we calculated five symptom factors^16^. See Table 1 for details of patients’ psychopathology. Healthy individuals who did not fulfill the criteria for any mental illness and were not in current or past psychotherapy of an ongoing mental health-related problem were recruited via online advertisement and flyers. None of the participants met the MRI exclusion criteria of claustrophobia, neurological disorders and metallic implants. Healthy individuals were recruited who matched the sample of patients with regard to age, sex, handedness and level of education (Table 1). Handedness was assessed with the Edinburgh Handedness Inventory (*n*=75), level of cognitive functioning was acquired with the Brief Assessment of Cognition in Schizophrenia^17^ (*n*=65) and verbal intelligence with a German Vocabulary Test^18^ (*n*=72). All procedures were approved by the ethics committee of the Charité-Universitätsmedizin Berlin.

**Table 1.** Demographics.

|  | Healthy<br>Participants | Schizophrenia<br>Patients | Statistics |  |
| --- | --- | --- | --- | --- |
|  | Mean (*SD*) | Mean (*SD*) | T (*DF*) | *P* Value |
| Sociodemographic Characteristics |  |  |  |  |
| Age (years) | 35.2 (*11.0*) | 35.3 (*10.8*) | -0.059 (*74*) | 0.953 |
| Gender | 24 male / 17 female | 21 male / 14 female |  |  |
| Edinburg Handedness Inventory<sup>a</sup> | 79.0 (*38.5*) | 81.47 (*36.3*) | -0.291 (*73*) | 0.772 |
| Education (years) | 14.1 (*2.9*) | 13.1 (*3.8*) | 1.345 (*74*) | 0.183 |
| BACS<sup>a</sup> | 270.4 (*37.0*) | 234.1 (*28.4*) | 4.427 (*63*) | < 0.001 |
| Verbal Intelligence (IQ)<sup>a</sup> | 100.5 (*10.6*) | 94.4 (*12.8*) | 2.204 (*70*) | 0.031 |
| Psychopathology |  |  |  |  |
| Illness duration (years) |  | 13.3 (*23.8*) |  |  |
| Illness onset (age in years) |  | 25.6 (*8.9*) |  |  |
| Chlorpromazine-equivalent (mg) |  | 317.1 (*221.6*) |  |  |
| PANSS Positive<sup>a</sup> |  | 16.9 (*8.6*) |  |  |
| PANSS Negative<sup>a</sup> |  | 16.6 (*8.1*) |  |  |
| PANSS Disorganized<sup>a</sup> |  | 21.1 (*7.9*) |  |  |
| PANSS Excitement<sup>a</sup> |  | 14.8 (*5.6*) |  |  |
| PANSS Distress<sup>a</sup> |  | 16.8 (*7.1*) |  |  |
| PANSS Total<sup>a</sup> |  | 63.6 (*20.0*) |  |  |
| SD, standard deviation; BACS, Brief Assessment of Cognition in Schizophrenia; PANSS, Positive and Negative Syndrome Scale; SAPS, Scale for Assessment of Positive Symptoms; SANS, Scale for Assessment of Negative Symptoms; |  |  |  |  |
| <sup>a</sup> sum score of items reported. |  |  |  |  |

### Data acquisition

Images were collected on a Siemens Tim Trio 3T scanner (Erlangen, Germany) with a 12-channel head coil. Structural images were obtained using a T1-weighted magnetization prepared gradient-echo sequence (MPRAGE) based on the ADNI protocol (TR=2500ms; TE=4.77ms; TI=1100ms, acquisition matrix=256×256×176; flip angle = 7°; 1×1×1mm^3^ voxel size). Whole brain functional resting state images during 5 minutes were collected using a T2*-weighted EPI sequence sensitive to BOLD contrast (TR=2000ms, TE=30ms, image matrix=64×64, FOV=216mm, flip angle=80°, slice thickness=3.0mm, distance factor=20%, voxel size 3×3×3mm^3^, 36 axial slices). Before resting state data acquisition was started, participants were in the scanner for about 10 minutes during which a localizer and the anatomical images were acquired so that subjects could get used to the scanner noise. During resting state data acquisition participants were asked to close their eyes and relax during data acquisition.

### Data preprocessing

The fist 5 images were discarded due to steady-state longitudinal magnetization. Firstly, slice timing was applied and then the data was realigned. Structural individual T1 images were coregistered to functional images followed by segmentation into gray matter, white matter, and cerebrospinal fluid. Data was then spatially normalized to the MNI template. To improve signal-to-noise ratio, a spatially smoothed with a 6-mm FWHM were done. Motion and signals from white matter and cerebrospinal fluid were regressed, followed by data filtering (0.01 – 1 Hz) to reduce physiological high-frequency respiratory and cardiac noise and low-frequency drift. Finally, the data was detrended. In addition, to control for motion, the voxel-specific mean framewise displacement (FD;^19^. FD values were below the default threshold of 0.5 for control and patient group (0.15±0.02 and 0.17±0.02, t-test p=0.42). All steps of data preprocessing were done using, SPM12 except filtering that was applied using REST toolbox^20^. SPM12 and REST toolbox were under MATLAB 2012b (www.mathworks.com).

### Network analyses

#### Functional connectivity matrices

In order to build functional networks (Fig. 1), nodes and links have to be defined and estimated. The nodes of the network were created based on the AAL atlas^21^. Nodes of which the averaged time series could not be estimated for all participants were excluded. The resulting network comprised 106 nodes. The node-averaged time series of BOLD signal were extracted for each subject using the REST toolbox^20^. Before the extraction, as is described in the data preprocessing section, the BOLD signal was corrected for motion, white matter and cerebrospinal fluid, all nuisance signals of no interest. The links between all 106 nodes, representing functional connectivity, were calculated using Person’s correlation coefficients (Fig. 1- (1,2,3)). Afterwards, a statistical threshold was applied assuring that only significant connections were taken to the next analysis steps. Non-significant links typically consisted of low correlations (< 10 %) and negative values.

**Figure 1.**
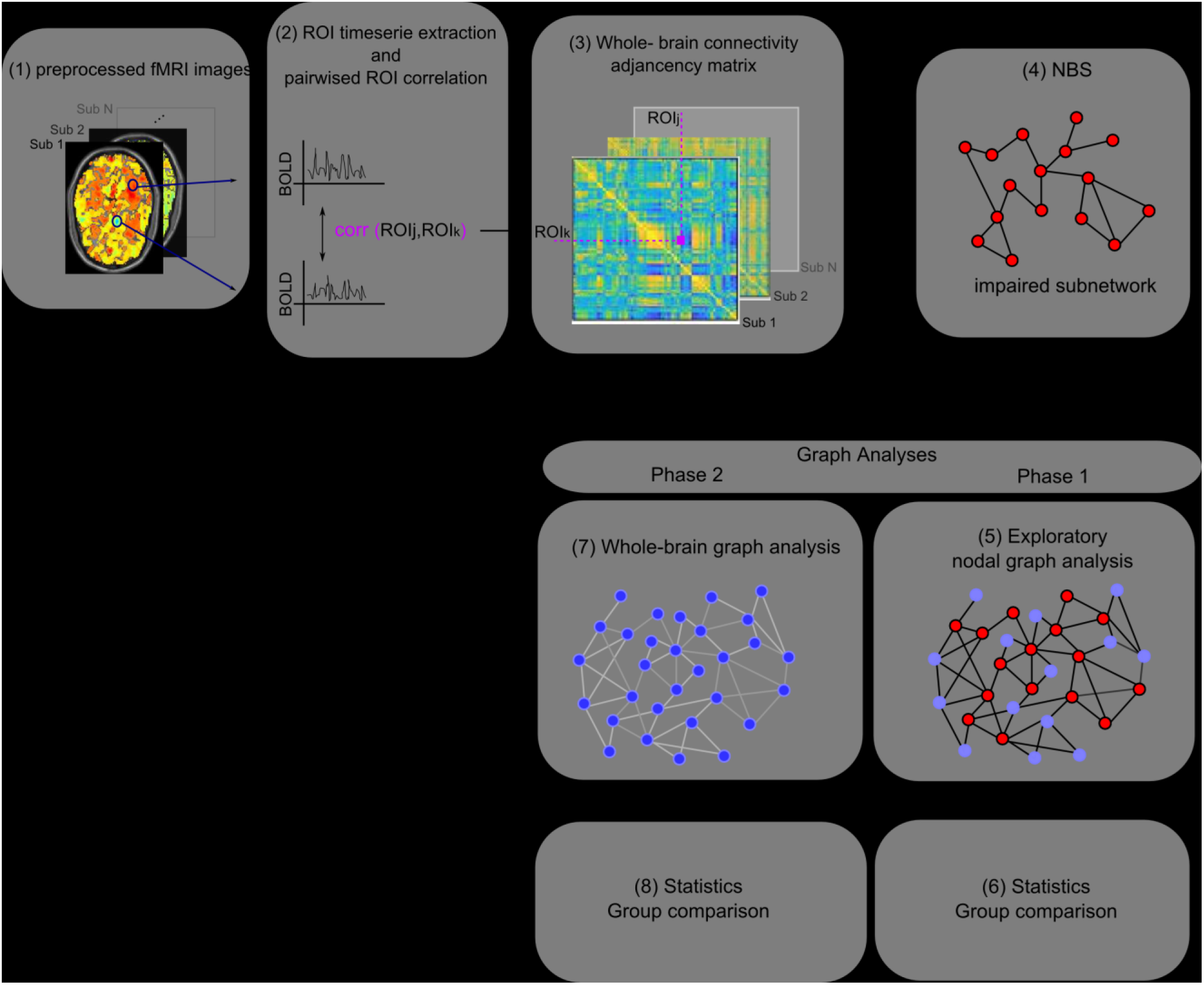
Analysis workflow. Resting state fMRI images were preprocessed (1) then the average timeseries of each ROI (AAL atlas) was extracted (2). Next, a pair-wise correlation of all timeseries using Pearson’s correlation coefficient forming the adjacency matrix (3). Network-based statistics was performed to extract the impaired subnetwork between groups (4). (5) Graph analysis Phase 1 – nodal graph analysis using nodes retrieved in (4). (6) Non parametric t-test of nodal graph measures calculated in (5). (7) Graph analysis Phase 2-whole brain graph analysis. (8) Non parametric t-test of graph measures calculated in (7).

#### Network-based statistics (NBS)

NBS is a nonparametric method to compare networks between groups based on cluster-based thresholding of statistical parametric maps. In contrast to other methods, instead of performing statistics in voxel clusters, NBS uses connected graph components, here represented by brain areas successive connected by links. These connected components are calculated from a set of suprathresholded links, using a breadth-first search algorithm. This procedure is repeated N times in order to estimate the null distribution. NBS results gain statistical power while controlling for family-wise error. The disadvantage of the method is the lack of specificity, as statistics are performed on connected graph components and thus no inferences can be made about individual links and nodes. For further details please refer to^14^ In this study, NBS was applied to functional connectivity networks (Fig. 1-4), calculated as described in functional connectivity matrices section.

#### Graph analysis

The topological properties of the functional networks were calculated using graph measures in the Brain Connectivity toolbox (brain-connectivity-toolbox.net)^22^. The following measures were considered: degree, which is the total number of connections to an individual node; betweenness that measures the fraction of all shortest paths that pass through an individual node; characteristic path length that measures the average shortest path between all pairs of nodes; efficiency that accounts for the average inverse shortest paths; diameter that is the longest of all shortest paths and cluster coefficient that is the fraction of a node’s neighbours that are also neighbours of each other. For a complete description of the graph measures please refer to^12,22^).

Graph analysis were conducted in 2 different phases: In phase 1 (Fig. 1- (5,6)), exploratory nodal analysis was conducted using nodes retrieved from the subnetwork in the NBS analysis to address the lack of specificity of the NBS method on the global range. In phase 2 (Fig.1- (7, 8)), graph analysis was applied to the whole brain network to characterize and compare the topology of the networks between schizophrenia patients and healthy subjects.

In order to build brain networks from the correlation matrices, a threshold is typically applied. Nevertheless, there is no consensus about the thresholding method, therefore it should be based on an educated guess^23^. Here, we used absolute thresholding ranging from 0 to 0.8 and weighted links as we believe that the strength of the connections play a role in schizophrenia.

### Statistics

#### NBS

NBS was run with N=10000 permutations to estimate the null distribution. The choice of the suprathreshold is an intrinsic and arbitrary step in NBS and it is known that high suprathreshold choices can omit components comprising the effect of interest^14^ Here we considered t statistic > 3 suprathresholds. The suprathreshold range in which significant group differences were found was 2.8 to 3.6 and we showed the subnetwork corresponding to suprathreshold 2.9 and t statistic > 3. To avoid potential confounding factors, the groups were fully matched for age, sex, handedness and level of education (Table 1) as detailed in the Participants section.

#### Graph measures

we used a nonparametric test namely random permutation. Permutations (N=10000) were performed to produce the null distribution of each measure where the p value was calculated and then thresholded using a significance level of p < 0.05. The multiple comparison problem was accounted by controlling the false discovery rate (FDR) at 5%. To avoid potential confounding factors, the groups were fully matched for age, sex, handedness and level of education (Table 1).

## 3. Results

### Network-based statistics (NBS)

Applying NBS (Fig. 1-4), we observed significantly increased functional connectivity (hyperconnectivity) in the group of schizophrenia patients compared to healthy individuals in a thalamo-cortico-cerebellar network consisting of the right thalamus, supplementary motor area, bilateral inferior occipital gyrus, bilateral superior occipital gyrus, right middle occipital gyrus, bilateral cuneus, bilateral lingual, right fusiform, left calcarine, left cerebellum III, vermis VII,VIII,IX (Fig. 2).

**Figure 2.**
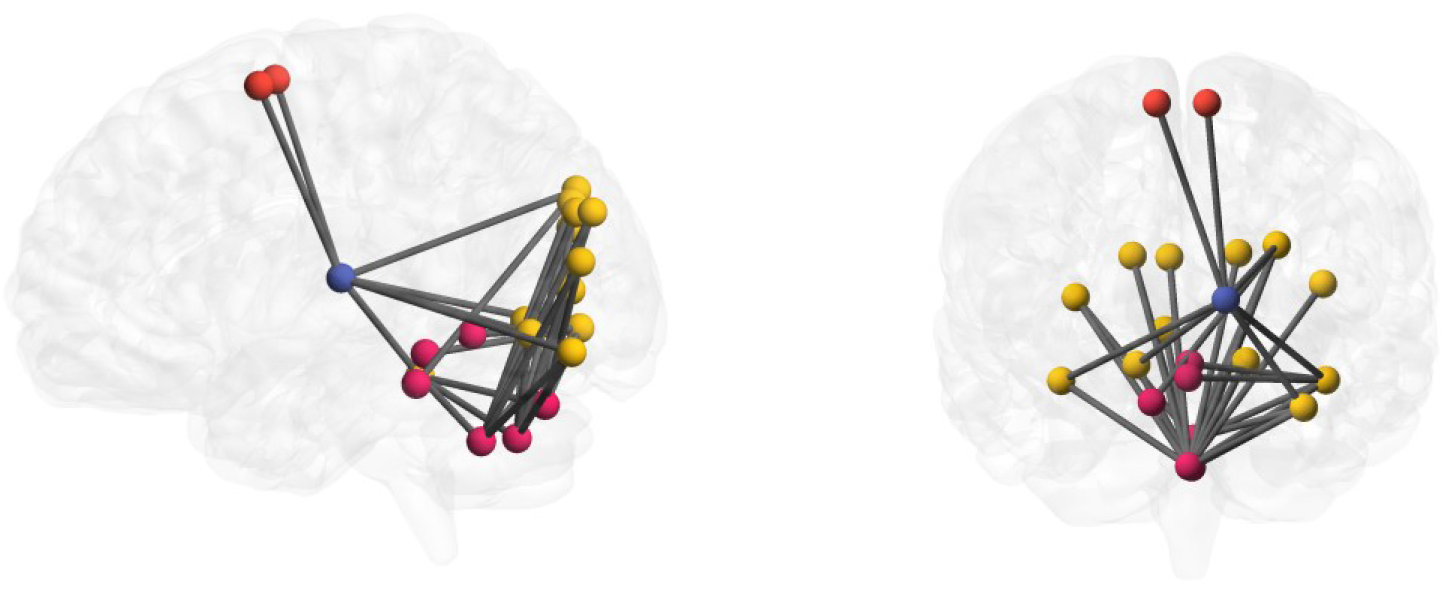
Subnetwork revealed by network-based statistic (NBS) analysis. The subnetwork showed alterations in a thalamo-cortico-cerebellar network comprising the thalamus (blue), superior motor area (red), superior, middle and inferior occipital gyrus (yellow), cuneus, lingual fusiform, calcarine, cerebellum III, vermis VII,VIII,IX (pink). Patients diagnosed with schizophrenia displayed significantly increased functional resting-state connectivity within this network compared to matched healthy individuals.

### Exploratory nodal graph analysis using nodes retrieved from the NBS subnetwork (Graph analysis Phase 1)

Due to the cluster-wise nature of the NBS, no inference can be made about single nodes and links from the impaired local subnetwork, resulting in a lack of specificity. Therefore, in order to explore the global role of the nodes belonging to the subnetwork revealed by NBS, we calculated their nodal topological characteristics in the whole brain network (Fig. 1- (5,6)): degree, betweenneess and cluster coefficient. Group difference across thresholds was found in the degree of thalamus with higher degree in patients with schizophrenia compared with controls (p=0.05 uncorrected).

### Whole brain graph networks analysis (Graph analysis Phase 2)

In order to investigate whether patients show alterations in global whole-brain topological properties of brain networks, we applied graph analysis to the whole brain functional connectivity network (Fig. 1- (7,8)). The measures considered were degree, betweenness, characteristic path length, global and local efficiency, diameter, and cluster coefficient. No group differences were found after correcting for multiple comparison.

## 4. Discussion

Schizophrenia is a complex mental illness characterized by a heterogeneous set of symptoms. Regarding its pathophysiology, different models have been proposed, which suggest that alterations in functional connectivity between different brain regions might underlie the variety of symptoms associated with schizophrenia^4,5^. Indeed, a growing body of empirical studies reported aberrant connectivity pattern in patients with schizophrenia where the integration of information across multiple brain areas is compromised resulting in disrupted brain networks^7–11,24^

In order to further explore the functional organization of brain networks in patients diagnosed with schizophrenia, we assessed local network differences using NBS and global network topological properties with graph analysis. NBS is a nonparametric method that directly compares functional networks between populations using connected structures from a suprathresholded set to identify subnetwork with abnormal connectivity among groups. Graph analysis is a popular tool to characterize the topology of networks, revealing central nodes of information processing as well as the network global informational capacity.

### NBS

The application of NBS on previously calculated functional connectivity networks revealed increased functional connectivity in a thalamico-cortico-cerebellar network in patients with schizophrenia compared to matched healthy individuals. This subnetwork consisted of the thalamus, supplementary motor area, temporal lobe, occipital lobe, and the cerebellum. The thalamo-cortico-cerebellar network has been assumed to play a key role in schizophrenia, since the thalamus is involved in regulating and integrating sensory information^25^ and can be thought as a gateway keeper or a filter. Andreasen^26^ hypothesized that symptoms characteristic for schizophrenia such as hallucinations, delusions, and self disturbances are due to an overload of information caused by a filtering disruption promoted by the thalamus^5,27^. However, it is not solely the thalamus that can explain the psychopathology of schizophrenia but rather its interaction with other areas such as the cerebellum^26^. Together they form the basis for cognitive dysmetria theory. Our result is in line with previous studies using diverse methods that indicated alterations in the functional connectivity of the thalamus and the cerebellum that will be discussed in the following. Regarding the thalamus, consistent with our finding is the frequent report of hyperconnectivity between the thalamus and sensory-motor regions^28–32^. Importantly, two of these studies performed brain-wide functional connectivity analysis on large sample sizes including more than 300 patients each^29,30^. Li et al.^30^ could additionally show that this pattern of thalamic hyperconnectivity is especially pronounced in chronic patients with schizophrenia, which is similar to our sample of chronic patients with a mean illness duration of 13 years. Second, regarding the cerebellum, previous studies have also shown altered functional connectivity of the cerebellum^33–35^. Although the function of the cerebellum is not yet completely understood, multiple models have been proposed in which the role of cerebellum in processing information from the motor cortex is extended to other cortical areas that are involved in higher order cognitive functions^8,36^. Multiple studies have shown structural alterations in this region in patients with schizophrenia including increases in cerebellar volume^37,38^, decreases in cerebellar volume^39–42^ and reductions in the gray matter that was associated with thought disorder^43^. Based on the assumed involvement of the cerebellum in a range of cognitive processes, Andreasen and Pierson^44^ hypothesized that pathological alterations within the cerebellum might have the potential to explain heterogeneous symptom characteristic for the diagnosis of schizophrenia. Moreover, from an information processing perspective, it has been suggested that a disconnection between cerebellum and cortex can lead to a misinterpretation of the information arriving from the cortex, resulting in, for example, experiences of delusion and auditory hallucinations^44^

Interestingly, when considering our finding of increased functional connectivity within the entire thalamo-cortico-cerebellar network, it also aligns with previous studies that showed alterations in thalamo-cortico-cerebellar circuits: increased connectivity between thalamus and multiple sensory areas, and decreased connectivity between thalamus and the cerebellum^32,45^, cerebellar region showed increased connectivity with the prefrontal cortex and thalamus as well as decreased connectivity with the visual cortex and sensorimotor cortex^35^, dysconnections in cognition-related resting state networks^46^, and reduced functional connectivity in a thalamo-cortico-cerebellar network^33,34,47^.

From the field of computational psychiatry further evidences for a role of the thalamo-cortico-cerebellar network in schizophrenia was obtained with cluster analysis of community detection^48^ and machine learning that predicted positive symptom progression^49^. Furthermore, the cerebellum was shown to have the highest discriminative power between patients and healthy controls^50^.

Concerning previous studies that used NBS in patients with schizophrenia patients and healthy controls showed wide-spread impaired connectivity. They are in line with our results when considering compromised heterogeneous multi-site communication in the brain in schizophrenia^51,52^,^14^,^53^. Specifically, the thalamus and the cerebellum were present in 2 studies showing structural disruptions in fronto-thalamic network^54^ and functional disruptions in the cerebellar-parietal and frontal areas^55^.

Taken together, multiple studies provided results that suggest an involvement of the thalamus in the psychopathology of schizophrenia. Notably, these results come from independent patient cohorts as well as methodologies, such as seed functional connectivity, whole brain connectivity, cluster analysis, independent component analysis and machine learning, which stresses its potential use as a biomarker.

### Exploratory nodal graph analysis

To cope with the lack of specificity of the NBS, we investigated the global role of the nodes belonging to the subnetwork revealed by NBS by calculating their nodal topological properties in the whole brain network. The results show higher degree in the thalamus in the group of patients when compared to healthy controls. As degree is a measure of centrality in the network that refers to the number of connections of a given node, it is related to the concept of a hub. Therefore, our finding indicates that the thalamus plays a central role in distributing information within the whole brain in patients with schizophrenia.

### Graph Analysis

We applied graph theory on the whole brain network, to explore global network differences. Interestingly, the whole brain graph analysis revealed no impairment in information processing on a global scale. Together with the NBS finding, the results of the current study suggest that altered connectivity is solely localized in the above-mentioned thalamo-cortico-cerebellar network in patients with schizophrenia. In line with this finding, a recent study^51^ also did not find group differences on a global scale. The authors along with Hadley (2016)^56^ proposed that clinically stable patients responding to medication regain network connectivity which therefore might become similar to that of the controls.

### Conclusion

Applying complex network analysis we observed a hyperconnected thalamo-cortico-cerebellar network in the group of patients diagnosed with schizophrenia suggesting that a greater amount of information is going through the thalamus reaching sensory areas and the cerebellum (and *vice-versa)* and that the thalamus thus plays a central role in the processing and distribution of information. Our findings emphasize that patients with schizophrenia suffer from multi-site communication problems in the brain and provide additional evidence for the involvement of the thalamo-cortical-cerebellar network in schizophrenia in line with the cognitive dysmetria model and the dysconnectivity theory. However, with the current status of the research it is not yet possible to determine whether these connectivity impairments are rather the cause or the consequence of a diagnosis of schizophrenia. To answer this important question, longitudinal studies over a long period of time, preferably containing data before the onset of illness are needed. Such studies will be also important for the development and effective testing of biomarkers.

